# Refolding activity of bacterial Hsp90 *in vivo* reveals ancient chaperoning function

**DOI:** 10.1101/462549

**Authors:** Tania Morán Luengo, Toveann Ahlnäs, Anna T. Hoekstra, Celia R. Berkers, Matthias P. Mayer, Stefan G. D. Rüdiger

## Abstract

The conserved molecular chaperones Hsp70 and Hsp90 play a key role in protein folding. Mechanistically, Hsp90 acts downstream from Hsp70 solving an Hsp70-inflicted folding block. It is unclear, though, when and to which extend the concerted action of this cascade becomes crucial in living organisms. Here we show that, in E. coli cells, Hsp90 dramatically improves protein refolding after heat stress while it is dispensable for de novo folding. We found that Hsp90 inhibition effectively reduced the refolding yields in vivo, leading to strongly reduced enzymatic activity of the paradigmatic chaperone client luciferase and broadly increased aggregation of the E. coli proteome. Additionally, the presence of Hsp90 during refolding reduces the net ATP consumption presumably by sparing the substrate binding-and-release cycles on Hsp70. This mechanism explains how the cooperation of Hsp90 with the Hsp70 chaperone system creates robust folding machinery in a sustainable manner. Together, we describe a general function for bacterial Hsp90 as a key factor of the folding cascade, which may be the ancient activity of this evolutionary conserved machine.

## INTRODUCTION

Molecular chaperones assist protein folding in the cell [1, 2]. The coordinated action of the conserved ATP-dependent Hsp70 and Hsp90 chaperone machines plays a key role for the attainment of the folded and mature condition of many proteins [3-6]. Hsp90 acts downstream of Hsp70, which together constitute an effective folding cascade [6]. This cascade acts early on the folding path, preparing their client for subsequent folding by itself.

A key step in this process is the transfer of the client from Hsp70 to Hsp90. This step stringently depends on the ATPase activity of Hsp90. [6-8]. Competitive inhibitors for ATP binding such as Geldanamycin and Radicicol act as potent conformational and functional Hsp90-specific blockers [9-12]. In eukaryotes, Hsp90 is involved in maturation and assembly of a wide array of protein substrates, many of which are involved in signalling processes [13, 14]. Recently, we have shown that Hsp90 also has a conserved key function in protein folding, solving an Hsp70-inflicted folding impasse [6]. Effective protein folding is crucial for all organisms, including bacteria. The bacterial Hsp90 system, HtpG, thus, offers the possibility to study when and under which circumstances the Hsp90 function in folding is crucial for the cell.

Here we show that *E. coli* Hsp90 plays a fundamental role in protein refolding after heat-shock in the cellular environment. Using the paradigmatic chaperone substrate firefly luciferase, we found that *in vivo* Hsp90 plays a key role in refolding after stress, while it is not essential for effective folding of nascent luciferase. The refolding function of Hsp90 is key to increase solubility of the global pool of cellular substrates after stress depending unfolding. The effectivity of the Hsp70-Hsp90 cascade also becomes evident when monitoring the ATP consumption, which we found to be 3-fold lower than for Hsp70 alone.

## RESULTS AND DISCUSSION

### Hsp90 is dispensable for *de novo* folding

To study the effect of Hsp90 in folding, we monitored bioluminescence of the model substrate luciferase from *Photinus pyralis* in living *E. coli* cells. This approach takes advantage of the fact that only fully folded luciferase is able to perform its enzymatic reaction. The reaction requires the firefly substrate D-luciferin, which is able to enter unpermeabilized *E. coli* cells and produces light that can be measured *in vivo* [15]. Luciferase is a monomeric multidomain protein, heterologous for *E. coli* and presents low refolding efficiency in the absence of chaperones, which makes it representative of many other substrates allowing generalisation [15, 16]. We confirmed that, in our hands, luciferase refolds *in vivo* after heat-induced denaturation in *E. coli* cells. We used an IPTG-inducible expression system to recombinantly produce luciferase in *E. coli* by induction before the stationary phase. We halted total translation by inhibiting the ribosome with tetracycline before temperature-induced unfolding of luciferase, ensuring the presence of a homogeneously unfolded luciferase population (**Fig. 1A**). After a 20-minute heat-shock at 42°C, when luciferase reached ~ 98% unfolding, we monitored the protein refolding over time. Luciferase produced in *E. coli* cells, containing both Hsp70 and Hsp90 wild type proteins, recovered up to ~60% after 45 min upon temperature downshift (**Fig. 1B**). This sets the scene for assaying the effect of Hsp90 in *de novo* folding and after thermal stress.

**Figure 1.**
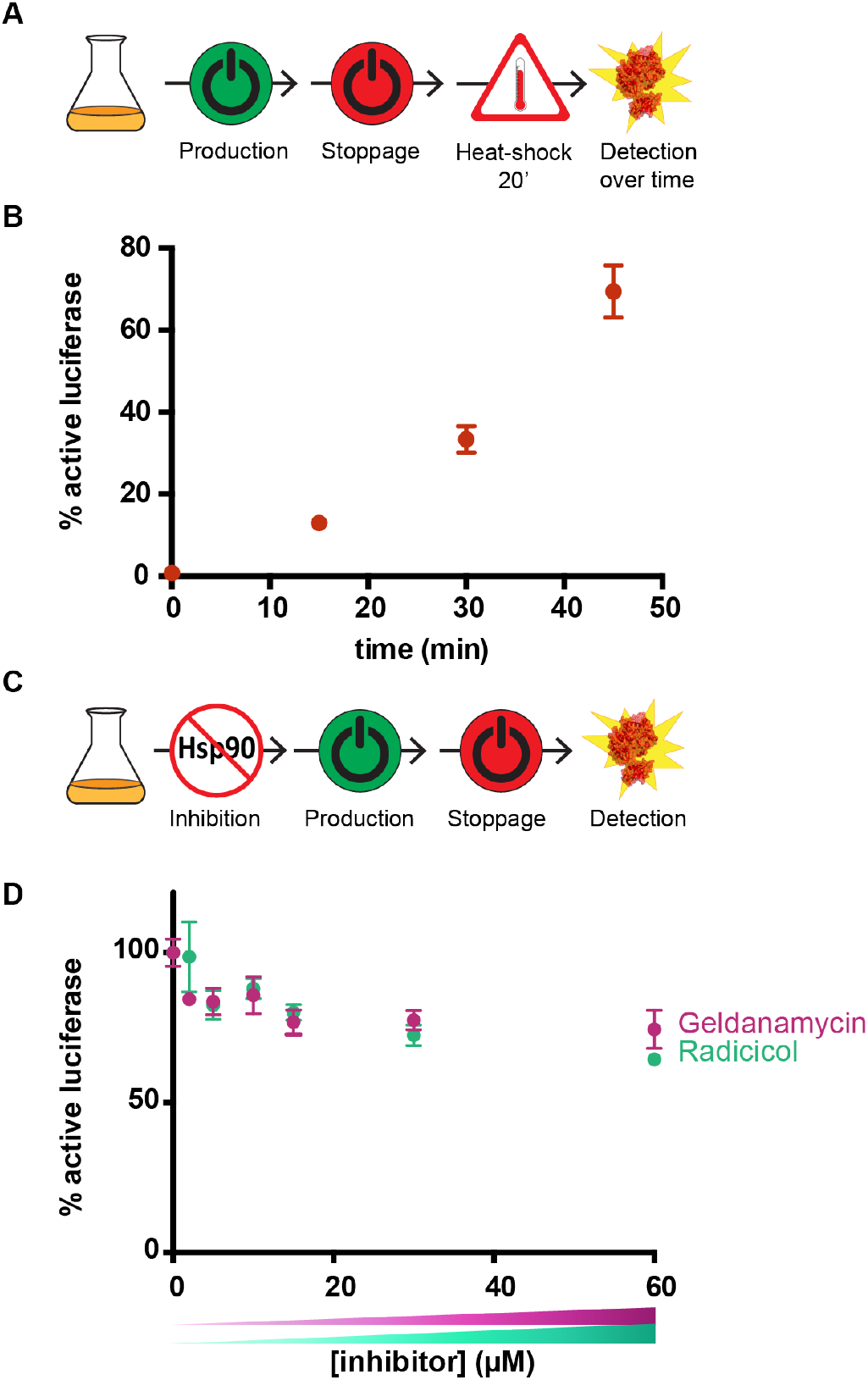
Hsp90 has only a mild influence in *de novo* protein folding. **A)** Experimental set-up for the luciferase refolding after heat-shock in *E. coli* shown in B). **B)** *In vivo* recovery of the luciferase activity after 20 minutes heat-induced denaturation. Luciferase recovery over time is plotted against the recovery time (±SEM n = 3). **C)** Experimental set up for the effect of Hsp90 inhibition in *de novo* folding. **D)** Geldanamycin and Radicicol have a weak effect in decreasing the production of active luciferase. Luciferase activity after production is plotted against increasing concentrations of the inhibitors (±SEM n = 3).

First, we tested the *in vivo* role of the bacterial Hsp90 for *de novo* folding. As Hsp90 function is tightly coupled to its ATPase activity, we used the Hsp90-specific ATPase inhibitors Geldanamycin and Radicicol to investigate the effect of Hsp90 in luciferase folding [10, 17, 18]. To assess the impact of Hsp90 inhibition in *de novo* protein folding, we blocked Hsp90 ATPase activity in *E. coli* cells by increasing concentrations of Geldanamycin and Radicicol (0 – 60 *μ*M) before production of luciferase allowing to assess the impact of Hsp90 inhibition in *de novo* protein folding (**Fig. 1C**). The amount of active luciferase did not decrease upon titration of each of the inhibitors (**Fig. 1D**). We conclude that Hsp90 is not essential for *de novo* folding of luciferase. This is remarkable as it is a paradigmatic chaperone substrate.

### Hsp90 plays a key role *in vivo* in refolding after heat stress

The function of Hsp90 is mechanistically important in protein refolding downstream from Hsp70 [6]. However, protein refolding and *de novo* folding strikingly differ in the availability of the folding domains, the protected environment created by the ribosome and the chaperones involved in the process [19]. In bacteria, newly synthesized proteins are welcomed by the ribosome-associated chaperone Trigger Factor (TF) [22]. TF and the Hsp70 bacterial homologue, DnaK, have combined activity and overlapping binding specificities [20, 21]. This implies that in the presence of TF, DnaK might not have access to the nascent chain and thus Hsp90 action is not required. Instead, upon stress conditions a big number of proteins are suddenly unfolded and prone to aggregate. In those situations, Hsp70 is essential for aggregation prevention and subsequent refolding [22]. To investigate the function of Hsp90 in this process downstream from Hsp70, we expressed luciferase in *E. coli* cells, and subsequently treated them with Geldanamycin or Radicicol (**Fig. 2A**). After 10 min incubation in the presence of the inhibitor, the cells were subjected to heat stress for 20 min at 42°C in order to achieve complete substrate unfolding, and we monitored the refolding of luciferase at different time points.

**Figure 2.**
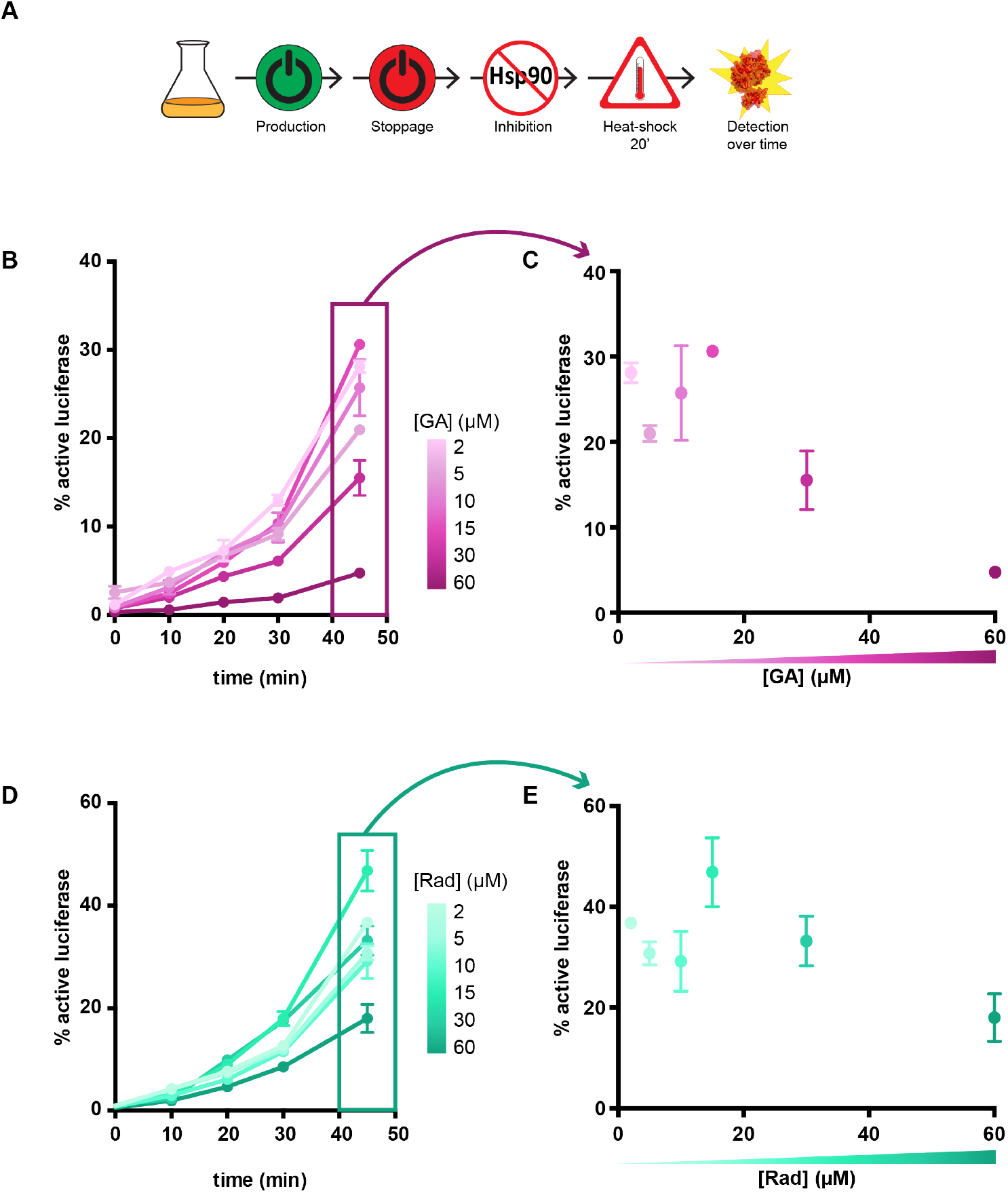
Hsp90 plays a key role in substrate activity recovery after heat-denaturation. **A)** Experimental set up for the inhibition of Hsp90 before heat-shock. **B)** Hsp90 inhibition with increasing concentrations of Geldanamycin impairs activity recovery. Luciferase activity is plotted against recovery time, increasingly intense colour representing increasing concentrations of Geldanamycin. **C)** The percentage of recovered luciferase after 45 min in B) is plotted against the concentration of Geldanamycin, showing a strong decrease of the recovery potential at the highest concentrations of the inhibitor used. **D)** Hsp90 inhibition with increasing concentrations of Radicicol impairs substrate activity recovery. Luciferase activity is plotted against recovery time, increasingly intense turquoise colour representing increasing concentrations of Radicicol. **E)** The percentage of recovered luciferase after 45 min in D) is plotted against the concentration of Radicicol, showing a decrease of the recovery potential at the highest concentrations of the inhibitor used.

Hsp90 inhibition by Geldanamycin causes a dramatic decrease in the refolding yield of luciferase (**Fig. 2B**). Increasing concentrations of Geldanamycin imply an eventual 6-fold reduction on the levels of active luciferase (**Fig. 2C**). A similar trend is observed when the Hsp90 inhibitor Radicicol was used (**Fig. 2D, E**). Both inhibitors specifically bind to the ATP pocket of Hsp90 although they do so with different affinities and different binding modes [11]. These results indicate that Hsp90 plays a key role in the *in vivo* refolding of luciferase after heat stress, in line with it being a stress-induced molecular chaperone.

### Bacterial Hsp90 is a general refolding chaperone

After establishing the role of Hsp90 in refolding for a single heterologous substrate, we went on to investigate its impact on the natural *E. coli* proteome. For this purpose, we used a solubility assay that allows distinction between soluble and higher molecular species. Unfolded or misfolded proteins have a higher tendency to aggregate so that monitoring the ratio of proteins in the pellet allows conclusions on the extent of the refolding reaction. The key advantage of this method is that it offers a fingerprint of the entire proteome of *E. coli*, which allows assessing how general the function of bacterial Hsp90 is. We compared the progression of refolding after heat stress in the absence and presence of Hsp90 inhibitor at different time points (**Fig. 3A**). By quantifying the level of proteins in the soluble and insoluble fraction after 45 min when refolding should be achieved for most proteins, we observe a striking increase in the protein levels found in the insoluble fraction when cells are treated with Geldanamycin (**Fig. 3B**). The recovery after heat-shock upon impaired Hsp90 occurred globally to a lesser extent after 45 min. Thus, bacterial Hsp90 plays an important and widespread ATP-dependent role on protein refolding after heat stress in the cell.

**Figure 3.**
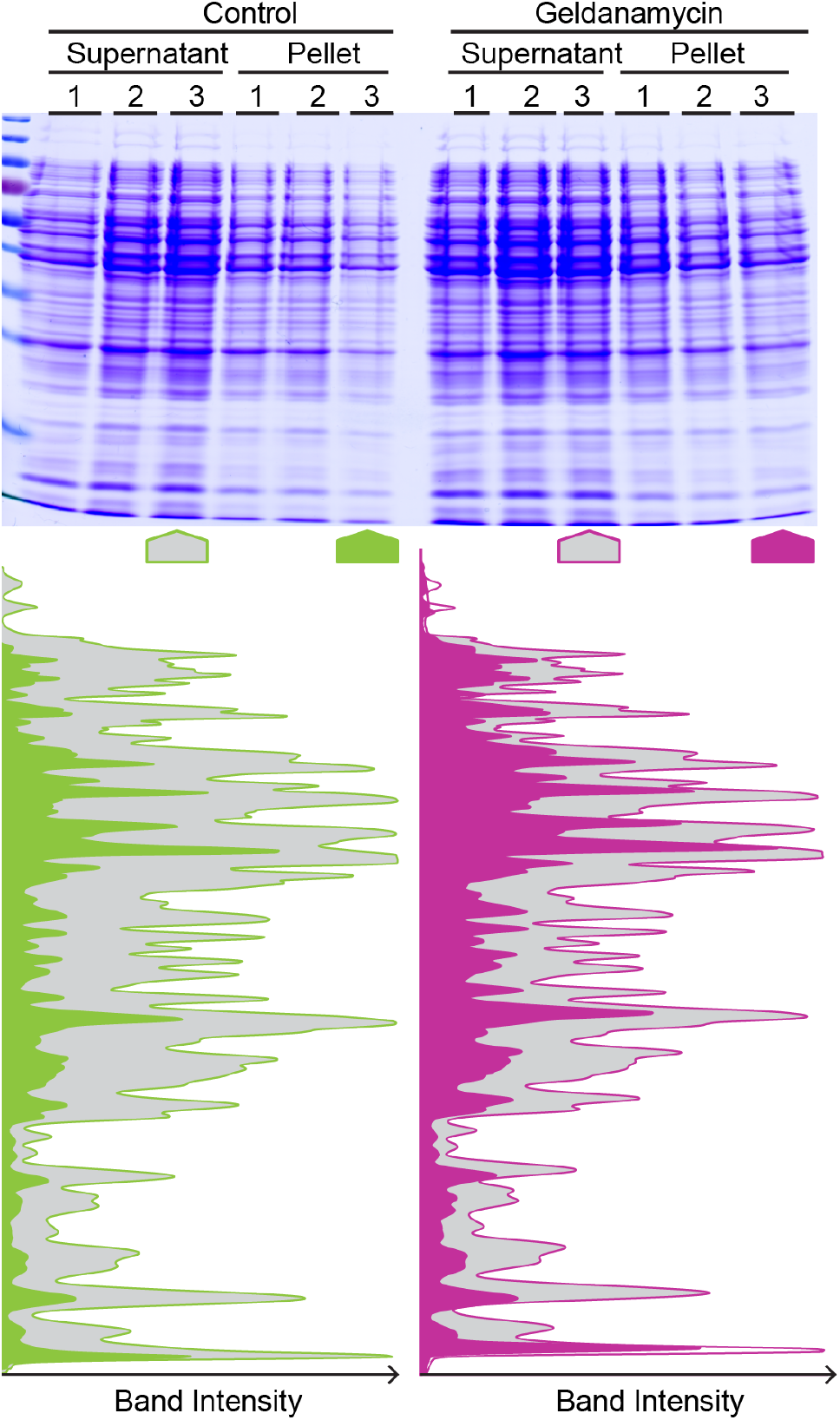
Hsp90 inhibition affects the general pool of substrates in the cytosol. **A)** Supernatant-pellet assay of total *E. coli* cellular pool in the presence and absence of the Hsp90 inhibitor Geldanamycin. Data points indicating (1) before heat-shock, (2) after heat-shock and (3) after 45 min recovery. **B)** Quantification of the SDS-page in A) after 45 min recovery. The graphs represent the intensity contrast between the supernatant band intensity (grey) and the pellet band intensity (green, control samples; magenta, geldanamycin treated samples).

### Hsp90 stabilises the native state

Next, we wondered whether Hsp90 may also influence the stability of a folded protein. Therefore, we looked at the unfolding pattern of the luciferase synthesized in the presence of increasing concentrations of the inhibitor geldanamycin. We applied a short heat-shock pulse of 3 min at 42°C, sufficient to unfold 50% of luciferase in the absence of inhibitor, and measured luciferase activity immediately after (**Fig. 4A**). We observe that high concentrations of the Hsp90 inhibitor lead to more severe luciferase unfolding, leaving only ~ 20% of active substrate (**Fig. 4B**). This might imply that, although the presence of the inhibitor did not cause a significant loss in the final enzymatic activity after translation, the produced luciferase is more sensitive to heat stress upon impaired Hsp90 activity.

**Figure 4.**
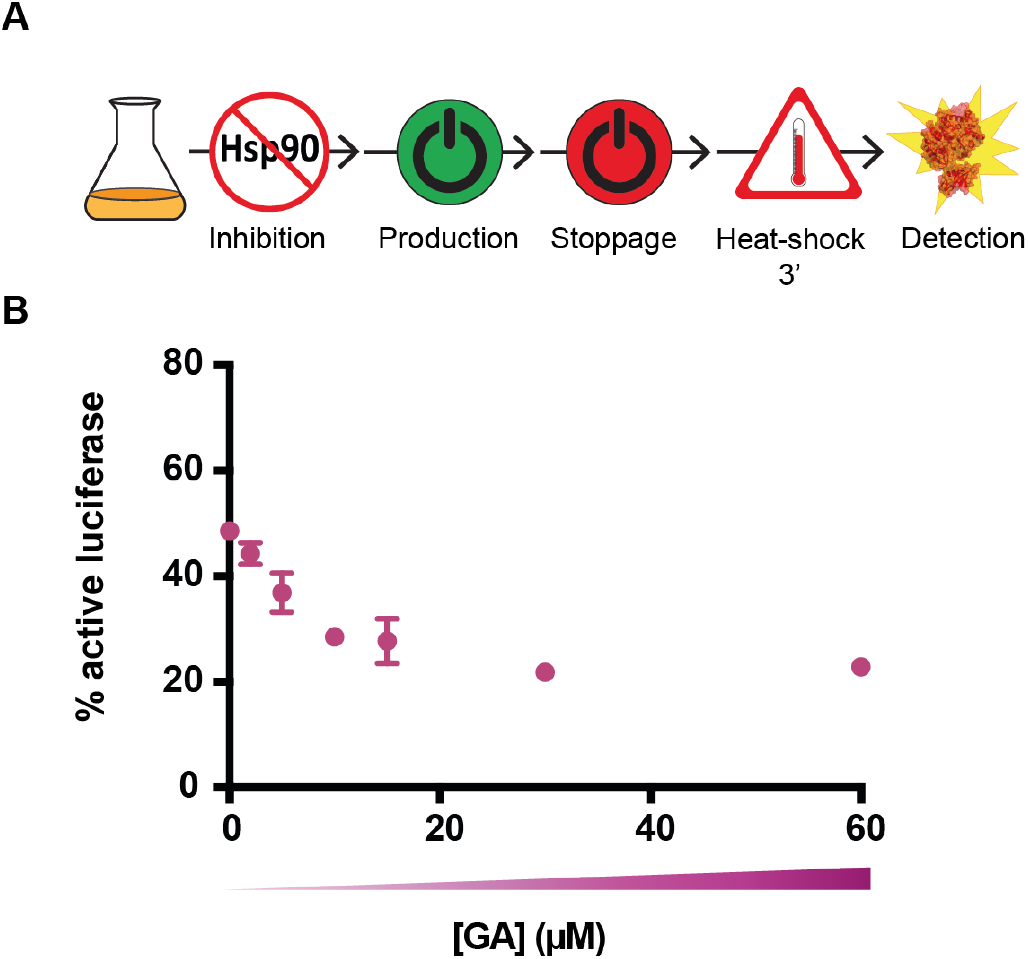
Hsp90 stabilises the native state. **A)** Experimental set up for the effect of Hsp90 inhibition by geldanamycin upon 3 min heat-shock. **B)** Geldanamycin inhibition renders the substrate more sensitive to brief heat-denaturation. At high GA concentrations, unfolding reaches lower yields after 3 minutes. Luciferase activity is plotted against the inhibitor concentration (±SEM, n = 3).

### Hsp90 reduces energy consumption for protein refolding

Our data establish that Hsp90 generally improves effectivity of posttranslational folding and stability. The functional and solubility assays demonstrate that Hsp90 plays an ATP-dependent role in protein refolding in the cell (**Fig. 2, 3**). This ATP-dependency is consistent with Hsp90 ATP hydrolysis being an essential requirement for substrate uptake from Hsp70 [6, 7]. In the absence of Hsp90, the Hsp70 system can promote folding under certain conditions by repetitive cycles of binding and release [6, 23, 24]. The presence of Hsp90 offers a straightforward path to the native state, which would reduce the number of Hsp70 cycles [6]. We hypothesize that the ability of Hsp90 to shortcut the path to the native state should overall reduce energy consumption of assisted folding.

To quantify ATP consumption during refolding, we used mass spectrometry-based metabolomics to monitor the production of ADP during protein folding *in vitro*. This allows linear quantification of ADP down to the μM level (**Fig. 5A**). We observed a 15-fold stimulation of the ATPase activity of the Hsp70 system by substoichiometric concentrations of the folding substrate luciferase after 30 min (**Fig. 5B**). This is in line with the synergistic ATPase activity stimulation by J-domain proteins together with substrates [25, 26]. Interestingly, the ATPase activity of Hsp70 did not decline over time although the refolding function of the chaperones is known to occur in the early stages of the folding reaction [6]. We suggest that, next to unproductive folding cycles, some aggregates might have formed during thermal unfolding keeping the Hsp70 system active and consuming ATP for a longer period of time. Strikingly, in the presence of *E. coli* Hsp90, HtpG, the total ATP consumption in the folding mixture is three-fold lower (**Fig. 5B**). This is an even more remarkable fact when considering that Hsp90 is an ATP driven machine itself, so the total ATPase activity would add up, if they worked independently. Mechanistically, this finding implies that Hsp90 reduces the number of Hsp70 binding-and-release cycles bringing down the total energy burden of refolding (**Fig. 5C**).

**Figure 5.**
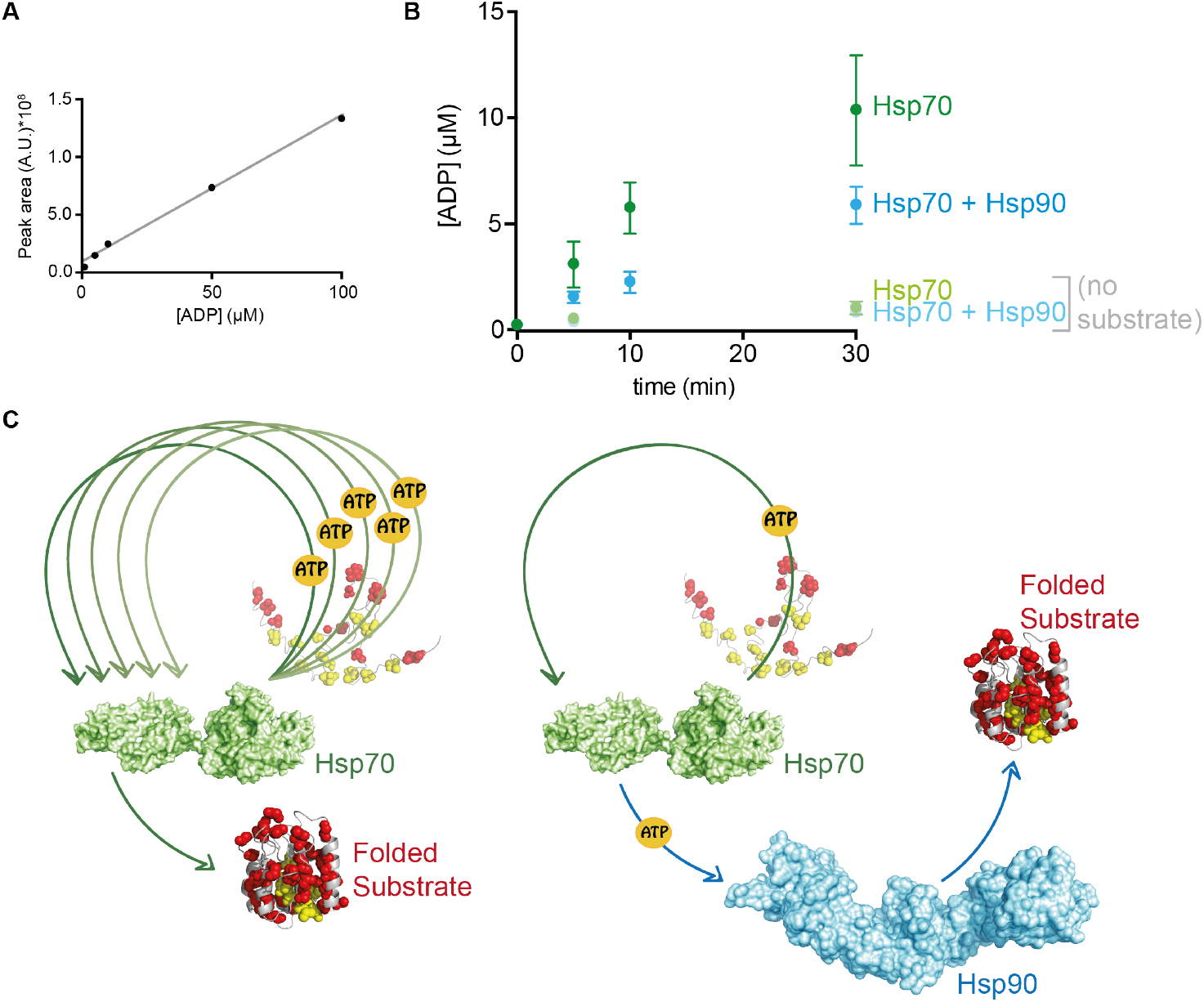
Hsp90 reduces substrate cycling on Hsp70 rendering folding energetically more efficient. **A)** ADP calibration curve showing the linearity of the detection range allows calculation of ADP produced in the refolding reaction. **B)** ADP production by the Hsp70 chaperone system – DnaK-DnaJ-GrpE– in the absence (green) and presence of HtpG (blue) determined by MS. Hsp90 minimises the production of ADP, and thus consumption of ATP over time. Data is shown as a result of three independent replicates ±SEM, n = 3. **C)** Proposed model for the energy usage optimization by Hsp90. Without the action of Hsp90, the unfolded substrate binds to Hsp70 and encounters consecutive cycles of binding and release, each of which hydrolyses an ATP molecule. In the presence of Hsp90, the substrate is transferred from Hsp70 to Hsp90 which also requires ATP but spares Hsp70 cycling yielding an energetically favourable process.

### Protein folding is assisted differently in depending on the situation

We analysed the role of Hsp90 for both *de novo* and posttranslational folding. We show that *E. coli* Hsp90 does not play a role during *de novo* protein folding. This is in agreement with previous findings for housekeeping Hsp70s and Hsp90s in eukaryotes [27, 28]. Thus, the Hsp70–Hsp90 chaperone cascade does not play a crucial role in the folding of nascent chains. Instead, upon protein biosynthesis, the nascent protein gets assistance of a plethora of other factors and ribosome associated proteins. In bacterial organisms, one of these factors is the chaperone Trigger Factor (TF) which is recruited shortly after the beginning of translation and binds the vast majority of nascent chains during translation [29, 30]. GroEL and DnaK are also able to bind non-native polypeptides generated by synthesis but in a posttranslational way and without interaction with the translational system [20, 31]. In eukaryotes, many more proteins fold and assemble co-translationally, and for that purpose there is a chaperone subset that is transcriptionally coregulated with the translational apparatus [32, 33]. In *E. coli* the Hsp70-Hsp90 chaperone cascade is dispensable in this process.

A special case of *de novo* folding in eukaryotes is the maturation process for signalling proteins that are yet to be activated. Regulatory proteins such as kinases and steroid hormone receptors involved in many signalling pathways require fine regulation by co-factors or post-translational modifications [14, 34]. Until then, they need to stay in a folding competent conformation, and this regulatory role is orchestrated by Hsp70 and especially, Hsp90 [7, 35]. Hsp90 function in maturation is, in turn, modulated by its plethora of co-chaperones [36]. Such a function is not needed for the bacterial Hsp90.

Instead, for posttranslational folding such as refolding after stress, bacterial Hsp90 has a key function *in vitro* and *in vivo* [6] (**Fig. 5C**). Here we show that Hsp90 is a key factor of this chaperone cascade despite the presence of countless substrates seeking chaperone assistance in the cellular environment. Hsp70 and Hsp90 directly interact in *E. coli*, and interestingly also in eukaryotic systems despite the presence of the linking co-chaperone Hop [37-40]. Hsp90 action comes after Hsp70 and the substrate transfer occurs in an ATP-dependent manner [6, 7].

### An efficient chaperone cascade for global unfolding

We show that the effective Hsp70–Hsp90 chaperone cascade has energetic implications, reducing the ATP cost of the folding process. Although Hsp70 is able to refold proteins on its own, binding-and-release cycles are energetically expensive. The folding protein needs to escape the tight binding of short and hydrophobic patches [41, 42]. Hsp90 offers a larger surface with many contacts distributed over the molecule that allow proteins to reach the native state [43, 44]. Key residues on this large surface sense the substrate and transmit the information to the other Hsp90 domains [45]. We propose a model in which the presence of Hsp90 spares re-cycling of the substrate on Hsp70, making the process more sustainable (**Fig. 5C**). This mode of action might be beneficial in particular after global stress events, both for effectivity of refolding and for its energy bill.

## MATERIALS AND METHODS

### In vivo luciferase refolding assay

#### Expression Control Experiment

XL1Gold E. coli cells containing the plasmid for Firefly Luciferase expression were grown overnight (37°C) in 50 ml of LB medium containing Ampicillin. The following morning, the culture was diluted to OD600=0.3, induced with IPTG (1 mM) and incubated at 20°C. After 30 min, tetracycline was added to a final concentration of 25 μg/ml in order to stop translation.

#### Hsp90EC de novo inhibition

XL1Gold E. coli cells containing the plasmid for Firefly Luciferase expression were grown overnight at 37°C in 50 ml of LB medium containing Ampicillin. The following morning, the culture was divided in several samples, diluted to OD600=0.3, and cells were incubated with different concentrations of the Hsp90 inhibitors Geldanamycin or Radicicol for 10 min, or DMSO as a control. Subsequently, cells were induced with IPTG to a final concentration of 1 mM and incubated at 20°C. After 30 min, tetracycline was added to a final concentration of 25 μg/ml in order to stop translation.

#### Hsp90EC inhibition before heat-stress

XL1Gold E. coli cells containing the plasmid for Firefly Luciferase expression were grown overnight at 37°C in 50 ml of LB medium containing Ampicillin. The following morning, the culture was divided in several samples, diluted to OD600=0.3, and induced with IPTG to a final concentration of 1 mM and incubated at 20°C. After 30 min, tetracycline was added to a final concentration of 25 *μ*g/ml in order to stop translation. Cells were then incubated with different concentrations of Geldanamycin, Radicicol or DMSO as a control for 10 min. After inhibition, cells were subjected to heat-shock for 20 min to 42°C.

#### Luminescence detection

Luciferase activity was measured in a luminometer plate reader (Berthold Technologies, XS3 LB930; MikroWin Software 2010). A sample volume of 10 *μ*l/well was added in a white, flat bottom 96-well plate (Corning Inc.), and 30 *μ*l of 100 μM luciferin substrate were injected by the instrument. Luciferin is able to enter non-permeabilized E. coli cells. Measurement was performed after 2 s delay with an integration time of 5 s. The activity of luciferase was normalized in respect to the control sample in the absence of inhibitors.

### Solubility assay

After Hsp90 inhibition and heat shock treatment and recovery, the 2 ml bacterial cultures were harvested by centrifugation (15 min, 6000 g). The bacterial pellet was resuspended in 30 *μ*l of 10x lysis buffer (100 mM Tris-HCl pH 7.5, 100 mM KCl, 2 mM EDTA, 15% sucrose) and lysozyme (1 mg/ml) was added at the moment of use. The resuspended pellet was stored at −80°C overnight. The following day, it was thawed on ice for two hours and 10- fold volume of ice-cold H2O was added and homogenized. Cells were lysed by sonication on ice (pulses: 5 min on/5 min off, 1 min). After sonication and a soft centrifugation step (15 min, 2000 g), 200 *μ*l of the supernatant fraction were removed to a new tube. The supernatant fraction was then spun down (15 min, 13000 g) after which, the supernatant was separated, and SDS-sample buffer was added to it. The pellet was resuspended in 600 *μ*l of lysis buffer with 2% NP40, sonication was applied again (pulse: 5 min on/5 min off, 30 s) and the sample was centrifuged (15’, 13000 g). The resulting pellet was resuspended in 50 *μ*l of SDS-sample buffer and both supernatant and pellet samples were unfolded and separated on a 12% SDS-gel followed by Coomassie Brilliant Blue staining.

### Protein Purification

E. coli DnaK was purified according to a published procedure [46]. Briefly, DnaK was purified as native protein with an N-terminal His6-Smt3-tag after overproduction in ΔdnaK52 cells (BB1994). Cell pellets were resuspended in lysis buffer (20 mM Tris/HCl pH 7.9, 100 mM KCl, 1 mM PMSF), subjected to lysis by microfluidizer EmulsiFlex C5 (Avestin, Ottawa, Canada), and afterwards applied onto a column with Ni-IDA resin (Macherey-Nagel, Düren, Germany). Subsequently, the column was washed with 20 CV of lysis buffer and 10 CV of ATP buffer (20 mM Tris/HCl pH 7.9, 100 mM KCl, 5mM MgCl2, 5 mM ATP), and 2 CV of lysis buffer. Proteins were eluted with elution buffer (20 mM Tris/HCl pH 7.9, 100 mM KCl, 250 mM imidazol). To remove the His6-Smt3 tag from DnaK, the protein was treated with Ulp1 protease. After cleavage and dialysis into lysis buffer, the protein mixture was subjected again to a Ni-IDA column and the flow-through fraction containing tag-free DnaK was collected. Subsequently, DnaK was bound to an anion exchange column (Resource™ Q, GE Healthcare) equilibrated in low salt buffer (40 mM HEPES/KOH pH 7.6, 100 mM KCl, 5 mM MgCl2). DnaK was eluted with a linear KCl gradient (0.1–1 M) within 10 CV.

E. coli DnaJ was purified according to a published procedure [47]. Briefly, DnaJ was purified as native protein after overproduction in the E. coli strain W3110. Cell pellets were resuspended in lysis buffer (50 mM Tris/HCl pH 8, 10 mM DTT, 0.6% (w/v) Brij 58, 1 mM PMSF, 0.8 g/l Lysozyme) and lysed by microfluidizer EmulsiFlex-C5. Cell debris was removed by centrifugation at 20,000 g for 30 min. One volume of buffer A (50 mM sodium phosphate buffer pH 7, 5 mM DTT, 1 mM EDTA, 0.1% (w/v) Brij 58) was added to the supernatant and DnaJ was precipitated by addition of (NH4)2SO4 to a final concentration of 65% (w/v). After centrifugation (15,000 g, 30 min), the ammonium sulfate pellet was dissolved in 220 ml buffer B (50 mM sodium phosphate buffer pH 7, 5 mM DTT, 1 mM EDTA, 0.1% (w/v) Brij 58, 2 M Urea) and dialysed against the 5 L buffer B. Subsequently, DnaJ was loaded onto a cation exchange column (SP-sepharose, equilibrated with buffer B), washed with buffer B and eluted with a 15 CV long linear gradient of 0 to 666 mM KCl. DnaJ containing fractions were pooled and dialysed against 5 l buffer C (50 mM Tris/HCl, pH 7.5, 2 M urea, 0.1% (w/v) Brij 58, 5 mM DTT, 50 mM KCl). Afterwards the sample was loaded onto a hydroxyapatite column equilibrated in buffer C. The column was first washed with 1 CV buffer C supplemented with 1 M KCl, and then with 2 CV of buffer C. DnaJ was eluted with a linear gradient (0-50%, 1 CV) of buffer D (50 mM Tris/HCl, pH 7.5, 2 M urea, 0.1% (w/v) Brij 58, 5 mM DTT, 50 mM KCl, 600 mM KH2PO4) and 2 CV of 50% buffer D. The DnaJ containing fractions were pooled and dialysed against 2 L buffer E (50 mM Tris/HCl, pH 7.7, 100 mM KCl).

E. coli GrpE was purified as described before [48]. Briefly, GrpE was purified after overproduction in ∆dnaK52 cells (BB1994). Upon expression, cell pellets were resuspended in lysis buffer (Tris/HCl 50 mM, pH 7.5, 100 mM KCl, 3 mM EDTA, 1 mM PMSF) and lysed using a microfluidizer EmulsiFlex-C5. The lysate was clarified by centrifugation (18,000 g, 50 min). To the cleared lysate, ammonium sulfate (0.35 g/ml) was added, and insoluble proteins were separated from soluble by centrifugation (10,000 g, 20 min). The pellet was dissolved in 200 ml buffer A (50 mM Tris/HCl pH 7.5, 100 mM KCl, 1 mM DTT, 1 mM EDTA, 10% glycerol) and dialysed twice against the same buffer (3 l, 4 h). Subsequently, protein was loaded onto an anion exchange column (HiTrap Q XL; GE Healthcare) equilibrated with buffer A and GrpE was eluted using a linear gradient of buffer B (50 mM Tris/HCl pH 7.5, 1 M KCl, 1 mM DTT, 1 mM EDTA, 10% glycerol). Fractions containing the protein were dialyzed against buffer C (10 mM KxHyPO4 pH 6.8, 1 mM DTT, 10% glycerol). On the following day the protein was loaded onto a Superdex 200 (GE Healthcare) gel filtration column equilibrated in buffer A and concentrated using HiTrap Q XL with a steep gradient.

E. coli HtpG was purified as a native protein after overproduction in MC1061 cells induced by L-arabinose. Upon expression, cell pellets were resuspended in lysis buffer (25 mM KxHyPO4 pH 7.2, protease inhibitor (cOmpleteTM, EDTA free, Roche) and 5 mM ß- mercaptoethanol). The cells were lysed by microfluidizer EmulsiFlex-C5 and the lysate was clarified by centrifugation (20,000 g, 40 min). The cleared lysate was loaded onto a Nickel column (Poros 20MC) and eluted with elution buffer (25 mM KxHyPO4 pH 8, 400 mM NaCl, 500 mM imidazole, 5 mM ß-mercaptoethanol). Eluted fractions were diluted in lysis buffer and loaded onto an anion exchange column (HiTrap Q XL). From this column, the protein was eluted using high salt buffer (50 mM KxHyPO4 pH 7.2, 1 M KCl, 1 mM ß-mercaptoethanol, 1 mM EDTA, 10% glycerol).

Photinus pyralis Firefly luciferase was purified according to previously described procedures [49]. Briefly, firefly luciferase was expressed in XL10 Gold^®^ strain (Stratagene, US). Cells containing the expression plasmid were grown at 37°C until OD600 = 0.5 was reached, at which point the temperature was lowered to 20°C. After 45 min shaking at 20°C, cells were induced with IPTG overnight. After harvesting by centrifugation, pellets were resuspended in precooled lysis buffer (50 mM NaxHyPO4 pH 8.0, 300 mM NaCl, 10 mM ß-mercaptoethanol, protease inhibitors (cOmpleteTM, EDTA free, Roche) DNase10 *μ*g/ml) and lysed using a microfluidizer EmulsiFlex-C5. The lysate was cleared and incubated with Ni-IDA resin for 30 min. Subsequently, the lysate-protino mixture was loaded onto a column and washed with 10 CV of lysis buffer, 10 CV of wash buffer (50 mM NaxHyPO4 pH 8.0, 300 mM NaCl and 10 mM β-mercaptoethanol) and eluted by addition of elution buffer (50 mM NaxHyPO4 pH 8.0, 300 mM NaCl, 250 mM Imidazole, 5 mM β- mercaptoethanol) collecting 1-2 ml fractions. Luciferase was dialyzed overnight using dialysis buffer (50 mM NaxHyPO4 pH 8.0, 300 mM NaCl and 10 mM β- mercaptoethanol, 10% glycerol).

### Determination of ATP consumption

The ATP activity of the chaperones in the refolding mix was calculated based on the ADP produced in the sample measured using LC-MS. Thermally denatured luciferase was diluted to a final concentration of 0.1 *μ*M into the refolding mix containing the indicated chaperones, in 200 mM NH4(OAc), 5 mM Mg(OAc)2, 5 mM β- mercaptoethanol and 2 mM ATP, pH=7.5. At the indicated timepoints, 1 μl of the reaction was diluted 100-fold in analysis buffer containing 40% methanol, 40% acetonitrile, 20% water. The analysis samples were shaken for 15 min at 4°C and centrifuged at 40000g for 15 min at 4°C. Supernatants were collected for LC-MS analysis. LC-MS analysis was performed on an Exactive mass spectrometer (Thermo Scientific) coupled to a Dionex Ultimate 3000 autosampler and pump (Thermo Scientific). The MS operated in polarity-switching mode with spray voltages of 4.5?kV and −3.5?kV. Metabolites were separated using a Sequant ZIC-pHILIC column (2.1 x 150 mm, 5 μm, guard column 2.1 x 20 mm, 5 μm; Merck) using a linear gradient of acetonitrile and eluent A (20 mM (NH4)2CO3, 0.1% NH4OH in ULC/MS grade water (Biosolve)). Flow rate was set at 150 μl/min. Metabolites were identified based on exact mass within 5 ppm and quantified based on peak areas of a calibration curve using LCquan™ software (Thermo Scientific). ADP calibration curve yielded the equation (peak area) = 1.264 * 106 * [ADP] + 8.216 * 106 (μM). Peak areas were within their respective linear range of detection and [ADP] was calculated adjusting the zero offset for the y axis.

### Data analysis and representation

Microsoft Excel and GraphPad Prism 6.0c software were used to calculate the mean and standard error of the mean. Data were plotted using GraphPad Prism 6.0c. The SDS-page results were quantified using ImageJ and data were plotted and coloured using Microsoft Excel and Adobe Illustrator CC 2018.

## ACKNOWLEDGEMENTS

We are grateful to Ineke Braakman for continuous support. We thank Madelon Maurice and Alba Cristobal for sharing instrumentation. We thank Esther A. Zaal for the support with the initial metabolomics test. S.G.D.R. was supported by Marie-Curie Actions of the 7th Framework programme of the EU [Innovative Doctoral Programme “ManiFold” (No. 317371) and Initial Training Network “WntsApp” (No. 608180)], the Internationale Stichting Alzheimer Onderzoek (ISAO; project “Chaperoning Tau Aggregation”; No. 14542) and a ZonMW TOP grant (“Chaperoning Axonal Transport in neurodegenerative disease”; No. 91215084).

## AUTHOR CONTRIBUTIONS

S.G.D.R., M.P.M. and T.M.L. conceived the study; S.G.D.R., C.R.B. and T.M.L. planned experiments; T.M.L., T.A. and A.T.H. performed the experiments; T.M.L. purified the proteins. T.M.L. analysed data; S.G.D.R. and T.M.L. wrote the manuscript, incorporating comments from all authors.

## REFERENCES

1. Saibil, H. (2013) Chaperone machines for protein folding, unfolding and disaggregation. Nat. Rev. Mol. Cell Biol. 14, 630–642

2. Kim, Y.E. et al. (2013) Molecular chaperone functions in protein folding and proteostasis. Annu. Rev. Biochem. 82, 323–355

3. Pratt, W.B. et al. (2008) The Hsp90 chaperone machinery regulates signaling by modulating ligand binding clefts. J. Biol. Chem. 283, 22885–22889

4. Taipale, M. et al. (2012) Quantitative analysis of HSP90-client interactions reveals principles of substrate recognition. Cell 150, 987–1001

5. Karagöz, G.E. and Rüdiger, S.G.D. (2015) Hsp90 interaction with clients. Trends Biochem. Sci. 40, 117–125

6. Morán Luengo, T. et al. (2018) Hsp90 Breaks the Deadlock of the Hsp70 Chaperone System. Mol. Cell 70, 545–552.e9

7. Kirschke, E. et al. (2014) Glucocorticoid receptor function regulated by coordinated action of the Hsp90 and Hsp70 chaperone cycles. Cell 157, 1685–1697

8. Mayer, M.P. and Le Breton, L. (2015) Hsp90: breaking the symmetry. Mol. Cell 58, 8–20

9. Whitesell, L. et al. (1994) Inhibition of heat shock protein HSP90-pp60v-src heteroprotein complex formation by benzoquinone ansamycins: essential role for stress proteins in oncogenic transformation. Proc. Natl. Acad. Sci. U. S. A. 91, 8324–8328

10. Grenert, J.P. et al. (1997) The amino-terminal domain of heat shock protein 90 (hsp90) that binds geldanamycin is an ATP/ADP switch domain that regulates hsp90 conformation. J Biol Chem 272, 23843–50

11. Roe, S.M. et al. (1999) Structural basis for inhibition of the Hsp90 molecular chaperone by the antitumor antibiotics radicicol and geldanamycin. J Med Chem 42, 260–6

12. Pearl, L.H. (2016) Review: The HSP90 molecular chaperone-an enigmatic ATPase. Biopolymers 105, 594–607

13. Taipale, M. et al. (2010) HSP90 at the hub of protein homeostasis: emerging mechanistic insights. Nature Reviews Molecular Cell Biology 11, 515–528

14. Schopf, F.H. et al. (2017) The HSP90 chaperone machinery. Nat. Rev. Mol. Cell Biol. 18, 345–360

15. Schröder, H. et al. (1993) DnaK, DnaJ and GrpE form a cellular chaperone machinery capable of repairing heat-induced protein damage. Embo j. 12, 4137–4144

16. Schumacher, R.J. et al. (1994) ATP-dependent chaperoning activity of reticulocyte lysate. The Journal of Biological Chemistry 269, 9493–9499

17. Prodromou, C. et al. (1997) Identification and structural characterization of the ATP/ADP-binding site in the Hsp90 molecular chaperone. Cell 90, 65–75

18. Panaretou, B. et al. (1998) ATP binding and hydrolysis are essential to the function of the Hsp90 molecular chaperone in vivo. Embo J 17, 4829–36

19. Evans, M.S. et al. (2008) Cotranslational folding promotes beta-helix formation and avoids aggregation in vivo. J. Mol. Biol. 383, 683–692

20. Deuerling, E. et al. (1999) Trigger factor and DnaK cooperate in folding of newly synthesized proteins. Nature 400, 693–696

21. Deuerling, E. et al. (2003) Trigger Factor and DnaK possess overlapping substrate pools and binding specificities. Mol. Microbiol. 47, 1317–1328

22. Hesterkamp, T. and Bukau, B. (1998) Role of the DnaK and HscA homologs of Hsp70 chaperones in protein folding in E.coli. Embo j. 17, 4818–4828

23. Ellis, J. (1987) Proteins as molecular chaperones. Nature 328, 378–379

24. Buchberger, A. et al. (2010) Protein quality control in the cytosol and the endoplasmic reticulum: brothers in arms. Mol. Cell 40, 238–252

25. Mayer, M.P. and Bukau, B. (2005) Hsp70 chaperones: cellular functions and molecular mechanism. Cell Mol. Life Sci. 62, 670–684

26. Liberek, K. et al. (1991) Escherichia coli DnaJ and GrpE heat shock proteins jointly stimulate ATPase activity of DnaKvProc. Natl. Acad. Sci. U. S. A. 88, 2874–2878

27. Nathan, D.F. et al. (1997) In vivo functions of the Saccharomyces cerevisiae Hsp90 chaperone. Proc. Natl. Acad. Sci. U. S. A. 94, 12949–12956

28. Svetlov, M.S. et al. (2006) Effective cotranslational folding of firefly luciferase without chaperones of the Hsp70 family. Protein Sci. 15, 242–247

29. Oh, E. et al. (2011) Selective ribosome profiling reveals the cotranslational chaperone action of trigger factor in vivo. Cell 147, 1295–1308

30. Becker, A.H. et al. (2013) Selective ribosome profiling as a tool for studying the interaction of chaperones and targeting factors with nascent polypeptide chains and ribosomes. Nat. Protoc. 8, 2212–2239

31. Teter, S.A. et al. (1999) Polypeptide flux through bacterial Hsp70: DnaK cooperates with trigger factor in chaperoning nascent chains. Cell 97, 755–765

32. Albanese, V. et al. (2006) Systems analyses reveal two chaperone networks with distinct functions in eukaryotic cells. Cell 124, 75–88

33. Shiber, A. et al. (2018) Cotranslational assembly of protein complexes in eukaryotes revealed by ribosome profiling. Nature 561, 268–272

34. Endicott, J.A. et al. (2012) The structural basis for control of eukaryotic protein kinases. Annu. Rev. Biochem. 81, 587–613

35. Verba, K.A. et al. (2016) Atomic structure of Hsp90-Cdc37-Cdk4 reveals that Hsp90 traps and stabilizes an unfolded kinase. Science 352, 1542–1547

36. Sahasrabudhe, P. et al. (2017) The Plasticity of the Hsp90 Co-chaperone System. Mol. Cell 67, 947–961.e5

37. Genest, O. et al. (2011) Heat shock protein 90 from Escherichia coli collaborates with the DnaK chaperone system in client protein remodeling. Proc. Natl. Acad. Sci. U. S. A. 108, 8206–8211

38. Alvira, S. et al. (2014) Structural characterization of the substrate transfer mechanism in Hsp70/Hsp90 folding machinery mediated by Hop. Nat. Commun. 5, 5484

39. Genest, O. et al. (2015) Hsp70 and Hsp90 of E. coli Directly Interact for Collaboration in Protein Remodeling. J. Mol. Biol. 427, 3877–3889

40. Kravats, A.N. et al. (2018) Functional and physical interaction between yeast Hsp90 and Hsp70. Proc. Natl. Acad. Sci. U. S. A.

41. Rüdiger, S. et al. (1997) Interaction of Hsp70 chaperones with substrates. Nat Struct Biol 4, 342–9

42. Mayer, M.P. (2013) Hsp70 chaperone dynamics and molecular mechanism. Trends Biochem. Sci. 38, 507–514

43. Karagöz, G.E. et al. (2014) Hsp90-Tau complex reveals molecular basis for specificity in chaperone action. Cell 156, 963–974

44. Radli, M. and Rüdiger, S.G.D. (2018) Dancing with the Diva: Hsp90-Client Interactions. J. Mol. Biol.

45. Rutz, D.A. et al. (2018) A switch point in the molecular chaperone Hsp90 responding to client interaction. Nat. Commun. 9, 1472–018-03946-x

46. Kityk, R. et al. (2015) Pathways of allosteric regulation in Hsp70 chaperones. Nat. Commun. 6, 8308

47. Graf, C. et al. (2009) Spatially and kinetically resolved changes in the conformational dynamics of the Hsp90 chaperone machine. Embo j. 28, 602–613

48. Schönfeld, H.J. et al. (1995) The DnaK chaperone system of Escherichia coli: quaternary structures and interactions of the DnaK and GrpE components. J. Biol. Chem. 270, 2183–2189

49. Rampelt, H. et al. (2012) Metazoan Hsp70 machines use Hsp110 to power protein disaggregation. Embo j. 31, 4221–4235

